# Reorganisation of Functional Connectivity Gradients in Post-Stroke Aphasia

**DOI:** 10.64898/2026.06.07.730675

**Authors:** Ramya Balakrishnan, Tirso Rene del Jesus Gonzalez Alam, Robert Leech, Cathy J Price, Elizabeth Jefferies

## Abstract

Aphasia after stroke arises from focal damage to language-relevant cortex but is accompanied by widespread alterations in functional connectivity. In 60 stroke survivors, we related principal component-derived language scores to both lesion location and stroke-related changes in resting-state functional organisation, combining structural lesion-symptom mapping with a gradient-based approach. Structural lesion-symptom mapping identifies which damaged regions are associated with specific language impairments but does not capture how surviving cortex is reconfigured within the brain’s large-scale architecture. In contrast, lesion-gradient mapping characterises whole-brain connectivity gradients – data-driven axes that summarise the principal patterns of variation in cortical connectivity – and quantifies how behavioural variation relates to displacement of structurally intact regions along these intrinsic axes. Stroke was associated with altered positioning of cortical regions along the gradient separating default mode and control networks. Notably, the position of the left inferior frontal gyrus along this axis predicted dissociable language outcomes: displacement toward default mode connectivity patterns was associated with better speech but poorer writing performance. This opposing behavioural profile suggests that inferior frontal cortex contributes to language through flexible large-scale coupling, with distinct coupling regimes differentially supporting spoken and written production. These findings indicate that gradient-based mapping provides a mechanistic link between focal tissue damage and distributed behavioural consequences by revealing systematic reorganisation of macroscale functional architecture beyond the lesion site.

## 1. Introduction

Aphasia is a common consequence of stroke, producing heterogeneous impairments across language tasks and other cognitive domains^1,2^. Patients may experience difficulties in comprehension, speech, reading, writing, and conversation, with major effects on quality of life^3,4^. This variability reflects the fact that language is supported by a distributed neural system spanning multiple cortical and subcortical regions^5^. Yet, because stroke lesions differ in their size and location, no two patients present with the same profile, making it challenging to predict outcomes and to tailor rehabilitation effectively. Lesion–symptom mapping has been instrumental in linking focal damage to specific deficits, identifying contributions of regions such as the anterior insula, inferior frontal gyrus, and precentral gyrus to speech fluency^6–8^. However, symptoms arise not only from damage to these areas, but also from disruptions to broader networks and disconnection between them^9–11^. Recovery is equally variable: some patients regain substantial function years after stroke, while others show persistent deficits^12–14^. Lesion location and size are important determinants of outcome, but they do not fully explain recovery trajectories, which, among other factors, also depend on functional reorganisation of surviving tissue and remote changes in whole-brain connectivity^15,16^.

Resting-state functional MRI has been widely used to characterise network-level changes after stroke, including connectivity between language regions, interhemispheric interactions between homotopic areas, data-driven identification of large-scale networks, and graph-theoretical measures of network properties such as modularity or hubness^17,18^. These methods have been valuable in showing that stroke alters both language-related and domain-general systems, including the default mode, salience, and dorsal attention networks^19,20^. However, each approach provides only a partial view: region-of-interest analyses are constrained by prior assumptions, interhemispheric measures capture only one dimension of connectivity, and data-driven techniques such as independent components analysis or graph theory fragment the system into discrete parts. What is missing is a holistic description that situates local and network-level changes within the brain’s continuous macroscale organisation ^21^. Such a perspective may better reveal how distributed functional systems support recovery and how focal lesions induce widespread alterations that shape cognitive and language outcomes.

Connectivity gradients offer a data-driven way to summarise large-scale patterns of functional connectivity across the cortex^21,22^. Rather than dividing the brain into discrete spatially defined networks, gradients position cortical regions along continuous axes based on similarities in their whole brain connectivity profiles, with each brain region occupying a unique position in an abstract gradient space. The principal connectivity gradient reflects the dominant pattern of variation in cortical connectivity and is commonly linked, post hoc, with unimodal sensory–motor connectivity at one pole and transmodal default mode regions at the other, capturing the hierarchy from local specialisation to global integration^21,23^. Additional higher-order gradients capture further, independent patterns of connectivity variation; for example, the second gradient differentiates between visual and somatomotor profiles, while the third distinguishes patterns typically associated with the control and default mode networks^24^. Critically, because gradients capture multiple independent components of connectivity, a region may appear normal along one axis but abnormal along another, reflecting different functional roles or task contexts. Language processing depends on coordination across distributed cortical regions including auditory areas for sound processing, temporal regions for word comprehension, and frontal regions for speech planning and production^25^. By situating each region within a continuous, multidimensional connectivity space, gradients provide a nuanced, component-specific characterisation of disrupted connectivity, revealing how stroke-related disruptions to functional connectivity relate to behavioural performance.

Here, we apply a combined lesion–symptom and lesion–gradient symptom mapping approach in a large cohort of stroke survivors from the PLORAS dataset who all had both structural and resting-state functional MRI as well as language and cognitive assessments. We asked three questions: (i) which lesion sites are associated with specific language deficits, (ii) how does stroke disrupt whole-brain functional connectivity as summarised by connectivity gradients, and (iii) how do gradient-based measures capture associations with language outcomes beyond focal lesion effects? By relating principal components of performance on the Comprehensive Aphasia Test to both lesion location and gradient-derived connectivity measures from resting-state fMRI, we aimed to move beyond focal accounts by incorporating large-scale patterns of functional connectivity gradients in the study of aphasia^26–28^.

## 2. Methods

### 2.1. Participants

The source dataset included 83 stroke survivors and 86 age-matched control participants from the PLORAS (Predict Language Outcome and Recovery After Stroke) dataset^29^. All participants were free from other neurological or psychiatric conditions such as dementia or depression, and were selected to have both a T1-weighted structural scan and resting-state fMRI. Stroke survivors completed an assessment of cognition and language using the Comprehensive Aphasia Test (CAT)^30^ and provided demographic information (age, gender, handedness, age at stroke onset, date of scanning and CAT). We applied further inclusion criteria to select participants with resting state fMRI scans suitable for functional connectivity analyses: i) a minimum of 70% valid scans; ii) no denoising failure cases; and iii) a mean head motion less than 0.35 mm.

The final sample consisted of 60 stroke survivors (24 females; mean age = 56.86 ± 10.74 years) and 79 age-matched controls (43 females; mean age = 38.23 ± 16.2 years). The mean age at stroke onset was 52.25 ± 11.51 years. All stroke participants were in the chronic phase, with a mean post-stroke duration of 55.31 ± 58.12 months. Among the 60 stroke participants, 36 had left-hemisphere lesions, 7 had right-hemisphere lesions, 15 had bilateral lesions, and 2 had lacunar infarcts. Lesion volume, estimated by the number of affected voxels, averaged 11,590 (SD = 11,320) voxels. Lesions spanned both cortical and subcortical regions, with the greatest overlap in the left superior corona radiata (present in 30 of 60 participants; Fig. 1.

**Fig. 1:**
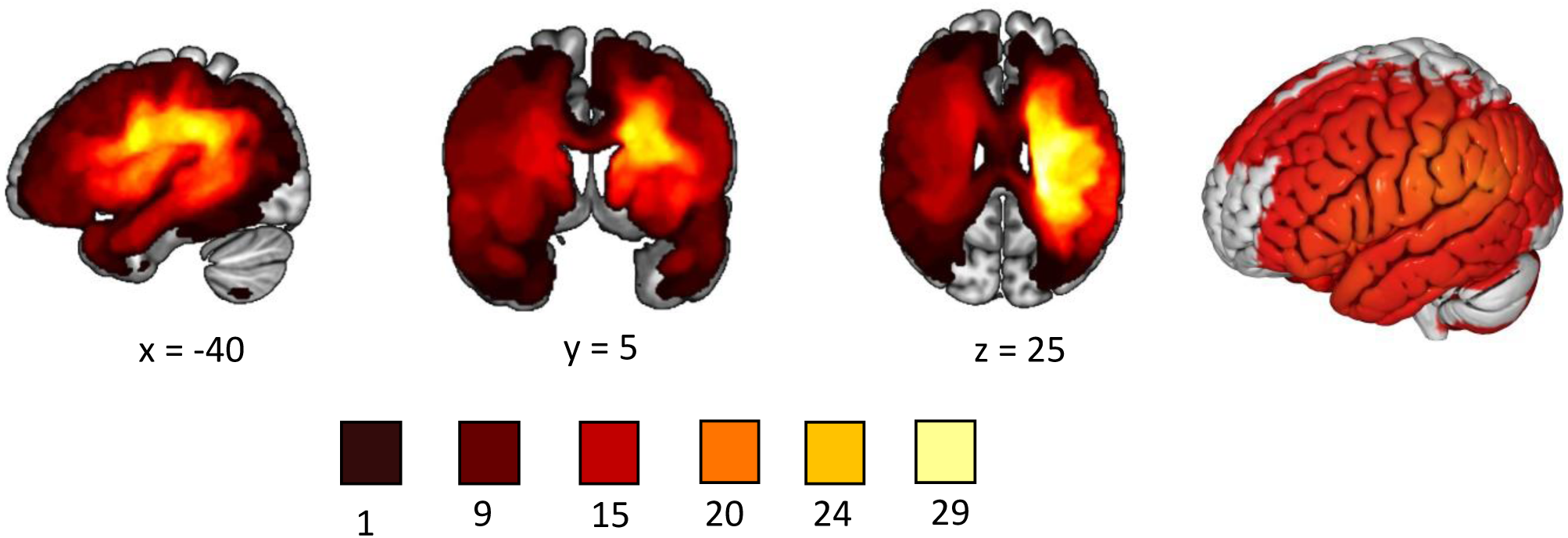
Overlap map of stroke lesions (N = 60). Binarised lesion maps were generated using FSLmaths^31^ and displayed in MNI space with MRIcroGL. Brighter colours indicate greater lesion overlap across stroke participants.

Across the sample, damage extended from frontal and parietal into occipital (visual) regions and portions of the temporal lobe, while sparing the temporal pole, temporal fusiform cortex, and parahippocampal gyrus.

### 2.2. Comprehensive aphasia test

Stroke-induced language dysfunction was examined using the Comprehensive Aphasia Test (CAT; Swinburn et al.,^30^). This test provides a broad assessment of aphasic patients’ performance in language functions and related cognitive domains. It includes language production tests (repetition, naming, and spoken picture description), as well as comprehension of spoken and written language, reading, writing, and repetition skills. Additional subtests evaluate cognitive domains, including line bisection, semantic memory, word fluency, recognition memory, gesture object use, and arithmetic.

Principal component analysis was performed to reduce these correlated language tests into distinct language domains explaining unique variance. First, we identified which subtests were suitable for inclusion by examining score variability using the interquartile range (IQR). Five subtests with few items and an IQR of zero, indicating that at least 50% of participants received the same score, were excluded (semantic memory, repetition of complex words, writing copy, comprehension of spoken paragraph, and reading functional words). PCA with oblique rotation was then performed on the standardised scores of the remaining subtests using R Studio. Components with eigenvalues greater than 1 were retained, yielding six components. The first four components loaded on multiple assessments and were readily interpretable as broader language domains; the remaining two components captured more task-specific variance and are reported in the Supplementary Materials (Section I).

### 2.3. Lesion segmentation

High-resolution whole-brain T1-weighted structural MRI (1 mm isotropic) was acquired for all participants, at the same time as the resting-state data, using research-dedicated 3T Siemens scanners at the Department of Imaging Neuroscience, UCL. Lesion segmentation was performed using established procedures implemented in SPM (running in MATLAB), in which each T1-weighted image was spatially normalised to MNI space and converted into a continuous lesion map that quantifies structural abnormality at each voxel on a scale from 0 (normal) to 1 (abnormal), relative to normative data from neurologically healthy controls^32^. Each lesion map was thresholded at 0.3 to generate binary lesion images. As this procedure has previously been shown to yield robust associations with behavioural outcomes, no other manual edits were applied. Segmented lesion maps were binarised using the fslmaths tool from FSL (FMRIB’s Software Library), and lesion–symptom mapping was performed on these maps.

### 2.4. Resting-state fMRI data preprocessing

We used the default resting-state fMRI preprocessing pipeline in CONN (https://www.nitrc.org/projects/conn)^33^, consisting of motion estimation, correction by volume realignment through a six-parameter rigid body transformation, slice time correction, and simultaneous white matter, grey matter, and cerebrospinal fluid segmentation and normalization to MNI152 stereotactic space (2 mm isotropic). These measures were applied to both functional and structural data. After preprocessing, the following potential confounding variables were statistically controlled for: six motion parameters determined at the previous step and their 1st- and 2nd-order derivatives; volumes with excessive movement (motion greater than 0.5 mm and global signal changes larger than z = 3); linear drifts; and five principal components of the signal from white matter and cerebrospinal fluid (CompCor approach)^34^. Then, a band-pass filter was applied to the data between 0.01 and 0.1 Hz. Global signal regression was not performed.

### 2.5. Whole-brain functional connectivity

After preprocessing and denoising the resting-state fMRI data for each participant in both groups (stroke and controls), we computed a functional connectivity (FC) matrix (Fig.2) and extracted the mean functional time series of each of the 400 cortical parcels, based on the Schaefer parcellation^35^ (https://github.com/ThomasYeoLab/CBIG/tree/master/stable_projects/brain_parcellation/Sch <u>aefer2018_LocalGlobal</u>). We then constructed a 400 × 400 FC matrix by calculating pairwise Pearson correlations between the mean time series of all parcels. These FC matrices were then averaged within each group to generate group-level connectivity matrices separately for stroke participants and age-matched controls. Finally, each Schaefer parcel contributing to the functional connectivity matrices was assigned to one of the seven canonical large-scale functional networks defined by Yeo et al.,^36^ according to maximum overlap between parcel and network. The Yeo networks^36^ were derived from group-level clustering of resting-state functional connectivity in healthy individuals and are labelled: (1) visual, (2) somatomotor, (3) dorsal attention, (4) ventral attention (or salience), (5) limbic, (6) frontoparietal control, and (7) default mode. We used these network assignments purely for interpretive reference. They did not influence the construction of functional connectivity matrices or the estimation of connectivity gradients.

**Fig. 2:**
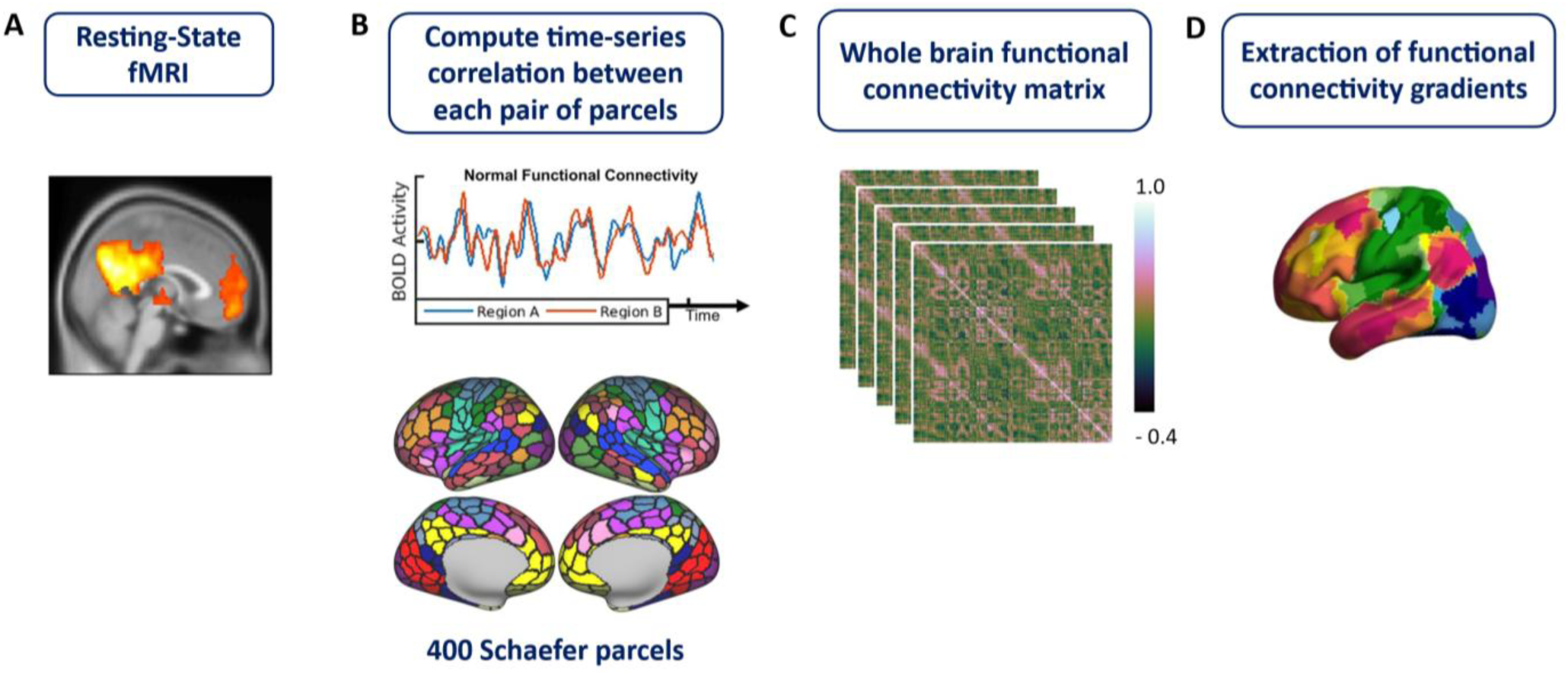
Processing pipeline for functional connectivity gradients in stroke participants and aged matched controls. A) Resting-state data of aged matched controls and stroke participants were pre-processed. B) Pearson correlations were computed between the mean time series of each of the 400 ROIs. C) A 400 × 400 functional connectivity matrix was generated for each participant in both groups. D) Functional connectivity gradients were extracted from the functional connectivity matrix for each participant.

### 2.6. Functional connectivity gradient extraction

Functional connectivity gradients for stroke participants and aged matched controls were extracted from the functional connectivity matrices using the BrainSpace Toolbox^37^. Healthy brains are expected to share common dominant axes of functional connectivity variation, with inter-subject variability in the precise positioning of regions along each axis. We focused on the first three gradients (Gradients 1-3), which capture the most dominant sources of variance in functional connectivity and have been linked to cognition in prior work^21,38^. To estimate Gradients 1–3 in each individual, we first extracted ten gradients from the functional connectivity matrix of each stroke participants and each age-matched control (dimensionality-reduction technique: diffusion embedding; kernel: normalized angle; sparsity: 0.9). The pipeline for gradient extraction is shown in Figure 2. Ten gradients were extracted because this allows the first three gradients to achieve a higher fit with canonical group-level gradients^21,39,40^. Gradients estimated from individual datasets can differ in orientation and ordering. Therefore, we aligned individual gradients for both stroke participants and controls to gradients extracted from the Human Connectome Project (HCP) dataset (n = 217; 122 women; mean ± SD age = 28.5 ± 3.7 years; Vos de Wael et al.,^41^) using Procrustes rotation. This procedure translates, rotates, and scales configurations to optimise alignment with the larger group average ^37^, while preserving the overall shape and structure of the data^42^. In other words, this alignment step improves interpretability by expressing individual gradients in a common coordinate system, facilitating comparison to gradients reported in other studies^21,39,40^.

### 2.7. Post-stroke gradient differences

Connectivity gradients for stroke participants contained effects of both stroke and aging. To isolate stroke-specific changes in connectivity, we subtracted each post-stroke gradient from the average gradient of age-matched controls included in this study using fslmaths (FMRIB’s Software Library). The resulting difference maps illustrate how each brain parcel shifted across the gradients following stroke; statistical analyses were then performed on these post-stroke gradient differences.

### 2.8. Statistical analyses

We conducted lesion–symptom mapping and lesion–gradient mapping. Lesion–gradient mapping was performed to identify probabilistic associations between lesion locations and the four PCA-derived components accounting for the largest sources of variance in language performance on the CAT. To directly compare lesion–symptom results with disruptions to gradient-based functional connectivity, we only examined lesions that damaged the cortex. This excluded 2 participants with lacunar infarcts and 6 with subcortical infarcts, leaving a sample of 52 stroke participants. Lesion–gradient symptom mapping addressed two questions: (1) What are the post-stroke changes in functional connectivity gradients? and (2) Can these changes in connectivity gradients predict language outcomes?

We used non-parametric t-tests in Randomise^43^ (https://fsl.fmrib.ox.ac.uk/fsl/fslwiki/Randomise/UserGuide), with 5,000 permutations, for both lesion–symptom and lesion–gradient mapping. For lesion–symptom mapping, the general linear model included binarised cortical lesion maps (n = 52) and each stroke participant’s language composite scores as covariates of interest, with whole-brain lesion size included as a covariate of no interest. Whole-brain lesion size was included to control for overall lesion burden, because behavioural outcomes reflect the total extent of the damage. Threshold-Free Cluster Enhancement (TFCE) was applied to identify significant, spatially extended lesioned clusters rather than relying solely on voxel-wise statistics^44^. Family-wise error correction was applied, and each contrast was thresholded at p < 0.05 to identify the brain regions significantly associated with each function.

For lesion–gradient mapping (n = 60), a general linear model was used to examine the association between stroke-induced connectivity-gradient changes (difference maps relative to healthy controls) and the four PCA-derived cognitive component scores. The model included the connectivity-gradient change maps and the four component scores as covariates of interest, with whole-brain lesion size entered as a covariate of no interest. Contrasts were set up for each PCA-derived component to identify parcels associated with poorer performance^45^. TFCE was implemented to identify parcels showing statistically significant associations between connectivity-gradient values and language outcomes, accounting for spatial contiguity across neighbouring parcels and correcting for multiple comparisons.

Lesion-gradient analyses were performed across the whole brain and within the Neurosynth-derived language-network mask. A lesion-overlap map was generated by combining cortical lesion masks from all participants. Voxels damaged in at least 25% of individuals were excluded to ensure analyses focused on connectivity changes in structurally intact tissue. This threshold corresponded to approximately 75% of all lesioned voxels, which were excluded from the whole-brain analysis. The same 25% threshold was applied to remove lesioned voxels from the language-network ROI. For visualisation, statistical maps for each contrast were thresholded at 0.95, corresponding to p < 0.05.

## 3. Results

### 3.1. Principal Component Analysis: Comprehensive Aphasia Test

The first four components accounted for 59% of the total variance, while two additional components explained a further 10%, giving a cumulative variance of 69%. The PCA loading matrix of CAT subtests and the scree plot illustrating the eigenvalues of each principal component are reported in Section I of the supplementary material. The first component (explaining 36% of the variance) had strong loadings on reading words (0.95), object naming (0.79), reading non-words (0.77), and reading complex words (0.75). The second component (explaining 9% of the variance) had strong loadings on repetition of digit strings (0.77) and sentence repetition (0.68) and was associated with phonology and working memory. The third component (explaining 7% of the variance) had high loadings for written picture naming (0.89) and writing to dictation (0.69) and was associated with writing and visuo-motor capacity. The fourth component (explaining 6% of the variance) showed strong loadings on comprehension of written words (0.82), comprehension of spoken words (0.54), and arithmetic (0.58), suggestive of semantic and broader cognitive difficulties, but was more mixed across tasks and showed more cross-loading with other components. Two further components had eigenvalues exceeding 1 but were dominated by loadings from single tasks assessing recognition memory (0.65) and gesture (0.94), respectively. As such, these components were treated as task-specific variance rather than coherent latent dimensions. They were included as covariates in subsequent brain analyses to account for this variance, but were not interpreted further (results are reported in Section I of the Supplementary Materials). PCA scores for each component were extracted; lower PCA scores reflected greater impairment on the respective component. These language component scores were used in lesion–symptom and lesion–gradient mapping analyses.

### 3.2. Lesion symptom mapping

Lesion–symptom mapping identified significant clusters associated with three language components from the PCA. PCA-2 (associated with poor phonology and working memory) showed the largest significant effect, which included left Heschl’s gyrus, parietal operculum, the anterior part of the superior temporal gyrus, the posterior part of the middle temporal gyrus, angular gyrus, supramarginal gyrus, and the superior division of the lateral occipital cortex (Fig. 3A). PCA-3 (associated with worse visuomotor performance) identified a cluster extending from the temporo-occipital part of the inferior temporal gyrus to the superior and inferior divisions of the lateral occipital cortex, as well as the superior parietal lobule, precuneus, and postcentral gyrus in the right hemisphere (Fig. 3B). PCA-4 (associated with poor performance on comprehension/arithmetic) identified the left middle temporal gyrus (temporo-occipital part) and the posterior part of the supramarginal gyrus (Fig. 3C). PCA-1 (associated with speech production) did not identify any significant effects in lesion-symptom mapping.

**Fig. 3:**
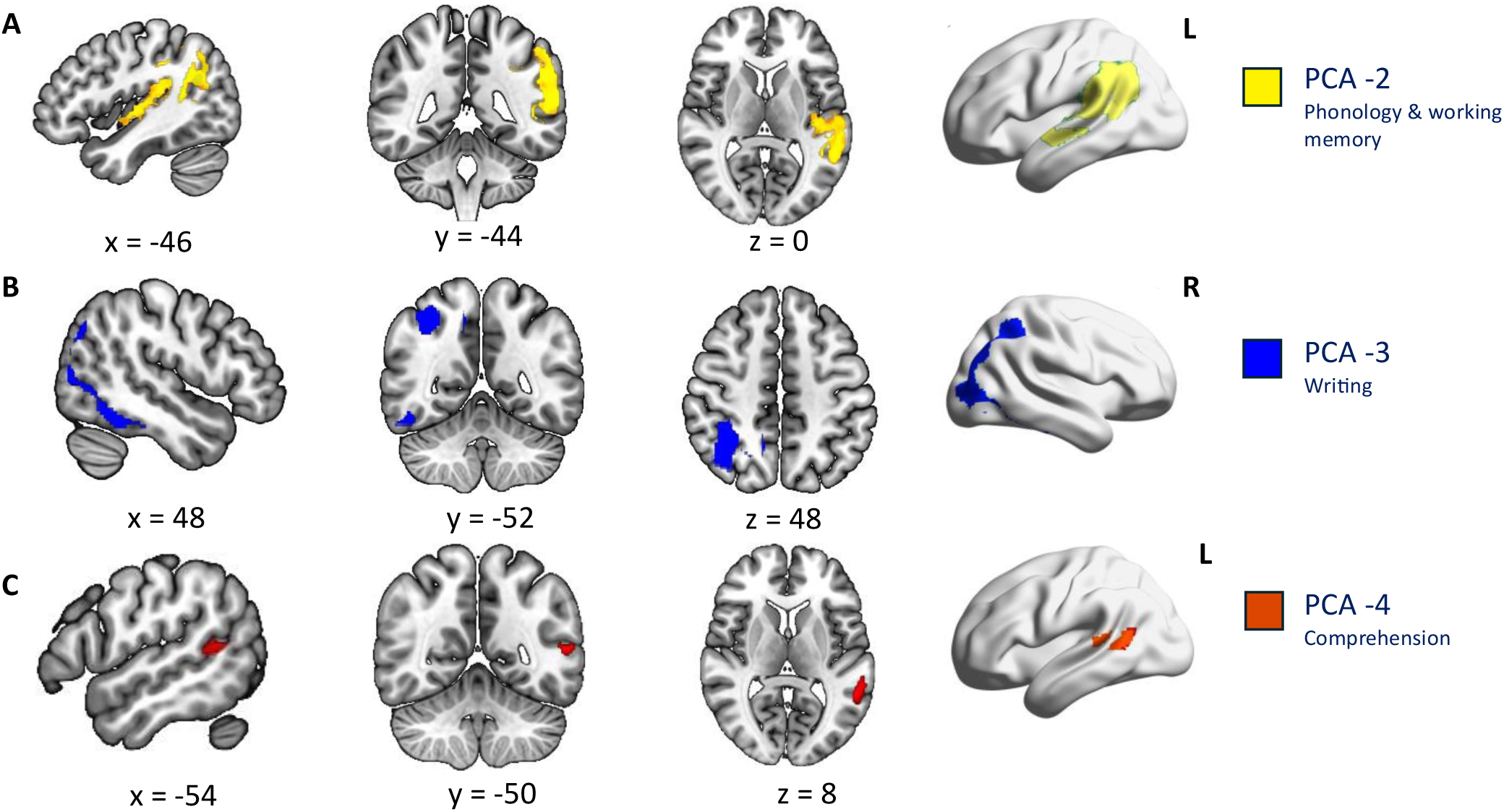
Lesion–symptom associations with PCA-derived language outcomes. A) Lesion locations associated with PCA-2 (phonology and working memory). B) Lesion locations associated with PCA-3 (writing and visuomotor capacity). C) Lesion locations associated with PCA-4 (comprehension/control difficulties).

### 3.3. Post-stroke gradient differences

Figure 4 shows connectivity gradients for young adults from the HCP dataset, plus the age-matched controls and stroke participants included in this study. Gradients for the young HCP sample, reproduced from Margulies et al.,^21^ are displayed using a warm-to-cool colour scale: warm colours indicate higher gradient values, and cool colours indicate lower values.

**Fig. 4:**
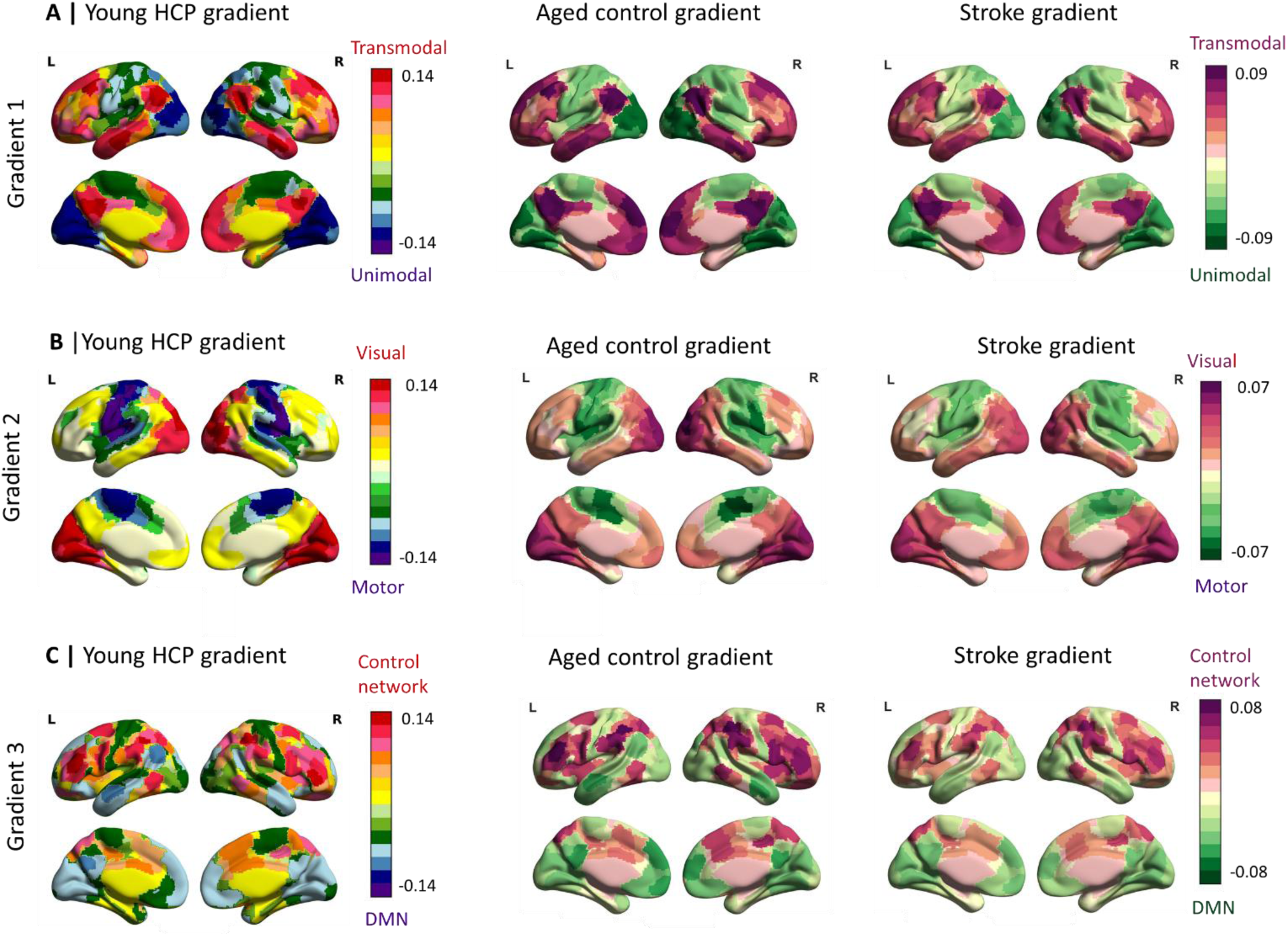
Functional connectivity gradients for age-matched controls and stroke participants from the PLORAS dataset, shown alongside canonical gradients from the young HCP sample (from Margulies et al.,^21^). In control and stroke participants, darker purple indicates higher gradient values and darker green indicates lower values. Panel A shows Gradient 1, representing the unimodal–to–transmodal axis. Panel B shows Gradient 2, representing the visual–to–motor axis. Panel C shows Gradient 3, distinguishing between the default mode and control networks. Compared with age-matched controls, stroke survivors show reduced gradient differentiation across all three gradients.

Gradients for the age-matched controls and stroke participants are shown using a purple-to-green colour scale, with darker purple indicating higher values and darker green indicating lower values. Gradient 1 differentiates unimodal (green) from transmodal (purple) regions; Gradient 2 separates motor (green) and visual (purple) regions; and Gradient 3 distinguishes default mode (green) and control networks (purple). Data for the stroke volunteers and controls are plotted on the same scale for each gradient. The maps are interpreted by comparing the colour assigned to each cortical region between these two groups; differences in colour reflect differences in a region’s position along the gradient.

While both the control and stroke participants show the basic gradient structure identified by Margulies et al.,^21^ in the HCP sample, some subtle differences in the magnitude of these gradients are visible – with less bright purple and green shades for the stroke survivors indicating that the magnitude of the connectivity gradients is attenuated following stroke (i.e. the networks that fall at opposite ends of these dimensions of connectivity are less distinct).

To isolate stroke-specific effects, we subtracted each stroke participant’s connectivity gradient from the average gradient map of age-matched controls, generating post-stroke difference maps for all three gradients. The average post-stroke difference maps for all three gradients are provided in Section II of the supplementary material. A non-parametric t-test with 5,000 permutations was used to determine significant post-stroke changes across the three gradients. Figure 5 shows the significant post-stroke changes on the gradients.

**Fig. 5.**
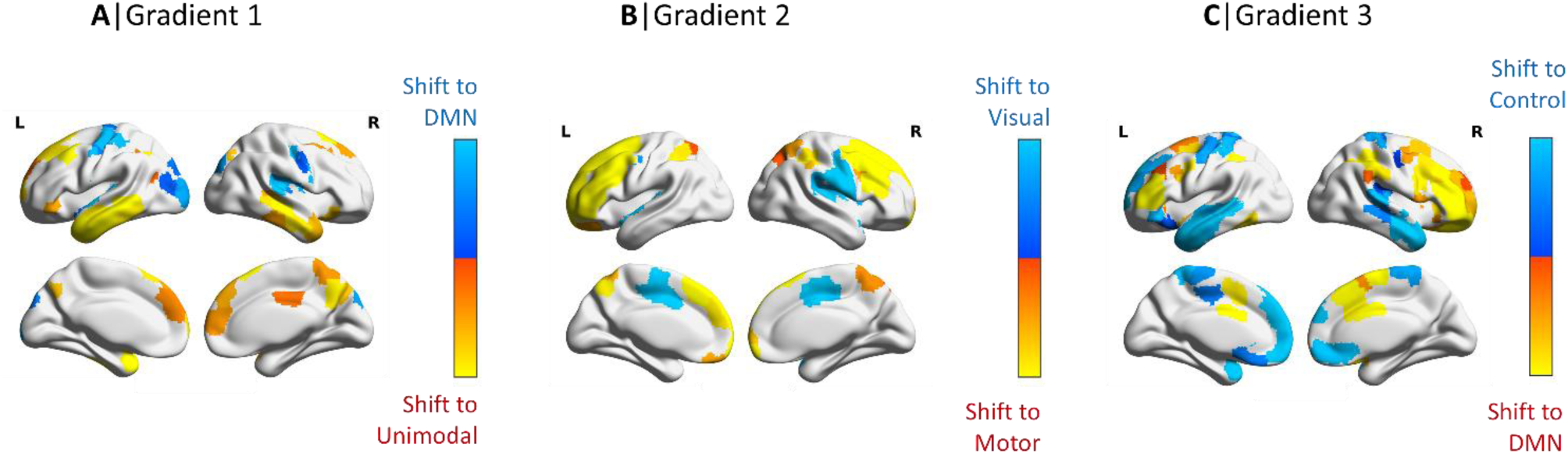
A–C. Significant post-stroke gradient changes for Gradients 1, 2, and 3 identified using permutation tests on gradient difference maps. Difference maps were generated by subtracting each stroke participant’s gradient map from the mean age-matched control gradient map. Warm-coloured regions indicate areas typically positioned toward the upper end of the gradient that shifted toward the lower end after stroke, whereas cool-coloured regions indicate areas typically positioned toward the lower end that shifted toward the upper end.

Gradient 1 (unimodal vs. transmodal) showed reduced differentiation after stroke, with transmodal regions—including the prefrontal cortex, temporal pole, middle and inferior temporal gyri, and right posterior cingulate—shifting toward the unimodal end of the gradient (Fig. 5A, warm colours). In contrast, unimodal regions such as the lateral occipital cortex and motor regions shifted toward the transmodal end (Fig. 5A, cool colours). Gradient 2 (visual vs. motor) also showed significant post-stroke changes: lateral and medial prefrontal regions shifted toward the motor end, while the pre- and postcentral gyri shifted toward the visual end, particularly in the right hemisphere (Fig. 5B). Gradient 3 (DMN vs. cognitive control) showed the most extensive post-stroke changes. Regions typically belonging to the control network—including the inferior frontal gyrus, frontal pole, supramarginal gyrus, and anterior cingulate cortex—shifted toward the DMN end of the spectrum (Fig. 5C). In contrast, DMN regions—including the medial prefrontal cortex, left dorsolateral prefrontal cortex, and lateral temporal cortex—as well as regions outside the DMN, such as motor areas, were positioned closer to the control-network end of this gradient axis (Fig. 5C).

### 3.4. Association between post stroke functional connectivity gradient changes and language outcomes within the language network

We examined how stroke-related changes in functional connectivity gradients within the language network related to PCA-derived language outcomes. We focussed on changes within the language network since the available behavioural data focussed on language assessments; changes in the position of language regions within the brain’s connectivity landscape are most likely to relate to these outcomes. To focus on intact tissue, voxels with ≥25% lesion involvement were excluded from the language-network mask (Fig. 6A).

**Fig. 6:**
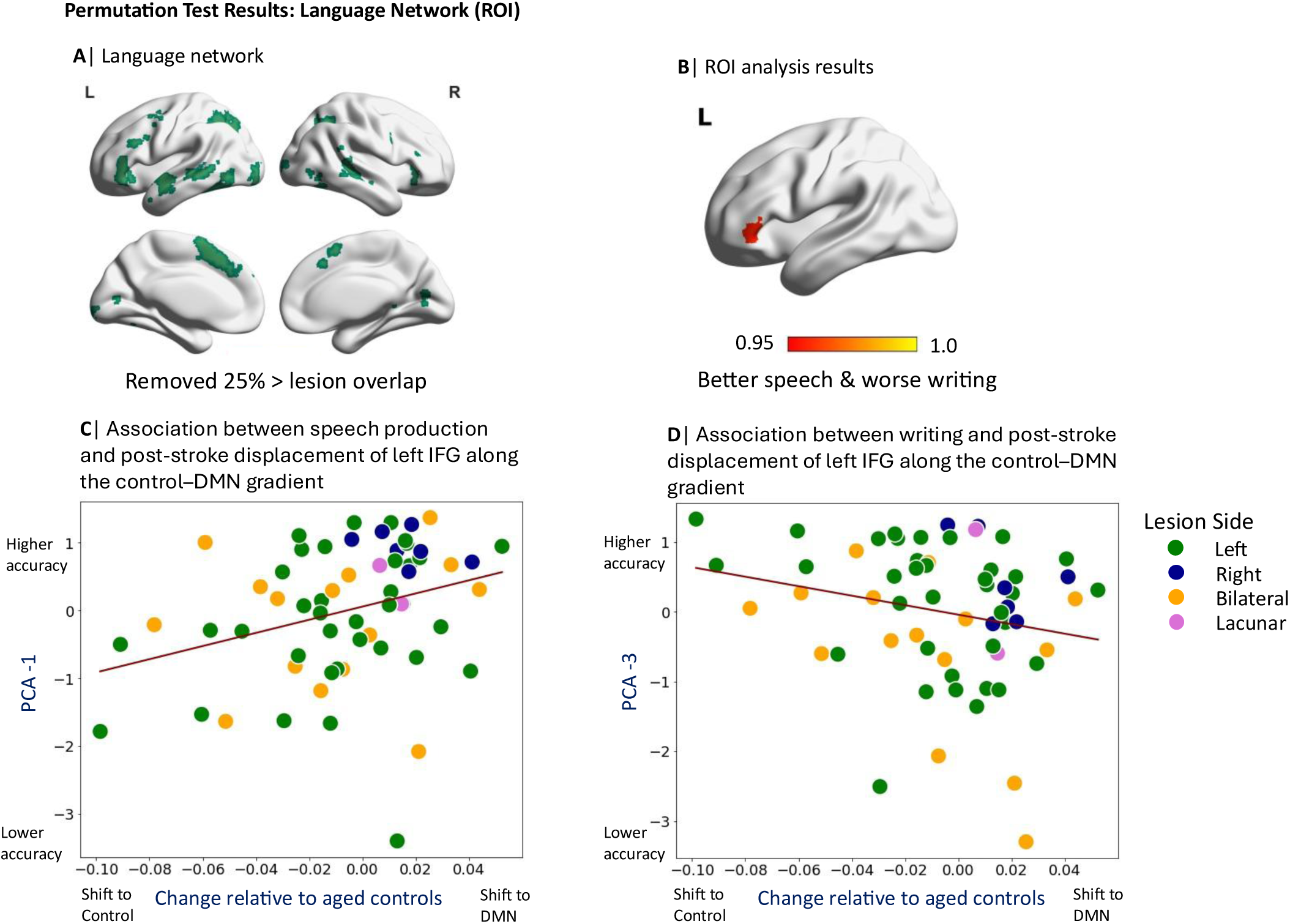
Permutation test results within the language ROI. A) Language-network mask used for the ROI analyses after removing lesion voxels. B) A significant cluster in the left inferior frontal gyrus (IFG) showed opposing behavioural associations along Gradient 3 (the default mode–control axis): displacement toward the default mode end (i.e. relatively greater DMN and reduced executive connectivity) was associated with higher PCA-1 scores (better speech production) and lower PCA-3 scores (poorer writing performance). C) Scatterplot showing the relationship between left IFG position along Gradient 3 and PCA-1 scores (speech production). D) Scatterplot showing the relationship between left IFG position along Gradient 3 and PCA-3 scores (writing ability).

Significant associations with Gradient 3 were found for PCA-1 (speech production), PCA-3 (writing), and PCA-5 (gesture–object use; reported in Section III of the Supplementary Materials, as PCA-5 loaded on a single task). Displacement of the left inferior frontal gyrus (IFG) pars triangularis (Fig. 5B) toward the DMN end of Gradient 3 was associated with better speech performance in stroke survivors (Fig. 5C). In healthy controls, this region is typically positioned toward the control network end of Gradient 3; thus, post-stroke movement toward the DMN reflects a relative shift in large-scale coupling. Notably, this same displacement of left IFG along Gradient 3 was associated with poorer writing to dictation (Fig. 5D), consistent with reduced engagement of control-related connectivity that may be required to coordinate linguistic, visuomotor, and executive processes during written output. These findings indicate that variation in IFG coupling along the DMN–control axis has dissociable behavioural consequences across language domains. A confirmatory permutation analysis found the results were unchanged when two participants who had lesions including this LIFG region were removed from the sample; PCA-1 and PCA-3 showed the same association with Gradient 3 displacement relative to age-matched controls.

To minimise the risk of Type II errors, we also conducted whole-brain lesion–gradient–symptom analyses relating gradient changes to PCA-derived language components, excluding voxels lesioned in ≥25% of participants. Full results are reported in Section IV of the Supplementary Materials.

## 4. Summary of results

Lesion–symptom mapping highlighted distinct neural substrates for specific language components: left temporoparietal damage was associated with verbal working memory deficits, right intraparietal sulcus lesions were linked to impaired visuo-motor tasks, and left posterior temporal lesions affected a component loading on comprehension and arithmetic, consistent with disruption to representational control processes. Beyond these structural effects, lesion–gradient mapping demonstrated that shifts in functional connectivity along the control–default mode axis (Gradient 3) captured cognitively meaningful aspects of functional reorganisation. Within the language network, left-hemispheric control regions exhibited greater functional affiliation with the DMN: a shift of the left inferior frontal gyrus towards this end of Gradient 3 predicted better speech production but poorer visuo-motor performance, reflecting domain-specific trade-offs in network reorganisation.

## 5. Discussion

This study shows that post-stroke language outcomes reflect both the location of structural damage and systematic variation in large-scale functional organisation captured by connectivity gradients. By integrating lesion–symptom mapping with gradient-based analyses, we provide a unified framework for understanding how focal injuries interact with the brain’s intrinsic macroscale architecture to shape behavioural variability after stroke.

Lesion mapping identifies regions in which tissue damage is associated with persistent language deficits, whereas gradient analyses quantify how surviving cortex is positioned along principal axes of functional organisation – particularly the axis separating default mode and control networks – and how displacement along these axes relates to language performance. Gradient displacement reflects structured reconfiguration of large-scale coupling patterns whose behavioural consequences depend on task demands. For example, variation in the position of left inferior frontal gyrus along the DMN–control axis was associated with opposing effects on speech and writing, indicating that differences in large-scale embedding of this region constrain distinct language functions. These findings situate post-stroke language variability within the geometry of intrinsic cortical organisation rather than attributing outcomes solely to lesion location.

### 5.1. Interpretation of lesion-symptom mapping

Lesion–symptom mapping was used to relate anatomical damage to behavioural impairment via PCA-derived dimensions of task performance. Our interpretation of these lesion–component associations draws on both the task loadings defining each component and prior evidence linking specific regions to cognitive functions; where these sources converge, the associations provide stronger inference about the functional consequences of focal damage. Convergent lesion evidence was observed for three PCA components (2, 3 and 4; see below). In contrast, no focal lesion effects were identified for PCA-1, which loaded heavily on speech production tasks. This absence of a localised lesion correlate suggests that variability in speech production in chronic aphasia may be less dependent on damage to a single cortical site and more influenced by distributed or reconfigured network dynamics. Consistent with this interpretation, PCA-1 showed significant associations with displacement along connectivity gradients (reported below), indicating that speech outcomes were better captured by variation in large-scale functional embedding than by focal tissue loss alone.

The strongest lesion effect was observed for the second PCA component. Behaviourally, this component loaded on tasks tapping phonological processing and verbal working memory. Anatomically, it was associated with damage to left posterior perisylvian and temporoparietal cortices. Similar lesions have been linked to repetition deficits in aphasia^46,47^, consistent with the role of the left superior temporal gyrus and supramarginal gyrus in phoneme identification^48,49^ and phonology^50,51^ and auditory short-term memory^52^.

The fourth PCA component was least specific, at the behavioural level capturing shared variance across spoken and written comprehension and arithmetic – perhaps indexing linguistic or representational control. However, anatomically, the associated lesion pattern involved the left posterior middle temporal gyrus, a region that integrates phonological information with lexical-semantic knowledge^53^ and acts as a convergence zone bridging DMN and multiple-demand control systems^54^. In addition, converging evidence has highlighted posterior temporal regions as critical for controlled semantic retrieval and regulation (e.g. Jefferies,^55^), providing a plausible anatomical context for this component despite its diffuse behavioural loadings.

Finally, the third PCA component weighted heavily on writing tasks, and less strongly on the line bisection and arithmetic tasks. The behavioural loadings therefore suggest a broader visuomotor function. It was associated with lesions extending into right occipital, parietal, and ventral temporal cortices. This pattern is consistent with a previous study linking right temporal lesions to writing deficits^56^ and aligns with accounts emphasising the reliance of writing on visual-motor integration^57–58^. Damage to the superior parietal lobule and precuneus are associated with visuospatial and working memory processes essential for orthographic planning^59^, whereas ventral temporal lesions impair semantic contributions to written output^60^. These findings reinforce the principle that persistent deficits on language assessments arise not only from injury to classical language regions but also from damage to multimodal hubs that support integration.

### 5.2. Reorganisation of gradient-based functional connectivity

Given that stroke often disrupts hub regions and white matter tracts supporting integration across networks^61^, functional reorganisation is expected to extend beyond damaged tissue. Connectivity gradients revealed stroke-related differences in large-scale functional organisation associated with language outcomes. Gradient 3 represents a continuum between control networks and the default mode network (DMN), providing a quantitative measure of how cortical regions are embedded within these systems. The left inferior frontal gyrus (IFG) pars triangularis occupied a relatively more DMN-weighted position along this axis in some individuals, reflecting comparatively stronger DMN and weaker control-network connectivity. Variation in IFG position along Gradient 3 was associated with better speech production but poorer writing and visuomotor performance, indicating that differences in its large-scale coupling profile relate differentially to distinct behavioural components.

Left IFG is widely regarded as a hub that coordinates distributed processes across cortical systems. Intracranial and neuroimaging evidence implicates this region in speech production^62^, and prior work suggests it occupies an intermediate position between control and default mode systems, enabling integration of conceptual representations with top-down control processes^54,63,64^. Within this framework, post-stroke variation in IFG embedding along the DMN–control axis can be interpreted as systematic alteration of large-scale coupling, with distinct consequences for different language and visuomotor demands

In healthy brains, the left IFG sits closer to the DMN than its right-hemisphere homologue, and this asymmetry relates to semantic selection ability^39^. In stroke survivors with aphasia, retaining or enhancing this DMN alignment may support better speech production by boosting word retrieval and selection mechanisms that underpin fluency^65,66^. However, the same shift toward the DMN was linked to more impaired writing and visuomotor performance, suggesting a functional trade-off in which tasks requiring stronger control-network engagement are disadvantaged^67^. Since the right IFG is typically positioned closer to control networks than the left IFG, one possible interpretation of our results is that shifts of left IFG toward the control end of this axis supports visuomotor demands at the expense of language fluency processes.

More broadly, the gradient changes observed within the language network are consistent with prior evidence that functional reorganisation in chronic stroke predominantly involves left-hemisphere language regions and, in some cases, their right-hemisphere homologues^68–70^. However, earlier studies largely relied on seed-based connectivity analyses, whereas the gradient framework characterises changes in the overall connectivity profile of language-network regions without requiring predefined seed regions. Rather than framing reorganisation as a binary shift between hemispheres, gradients quantify variation along continuous axes of large-scale organisation, revealing systematic alterations in how language regions are embedded within distributed networks. Such variation in large-scale embedding may reflect underlying plasticity processes, including stabilisation of surviving circuits and strengthening of alternative functional pathways when pre-existing networks no longer fully support language function^71^.

### 5.3. Integrating lesion location and gradient organisation

Taken together, lesion mapping identifies regions in which structural damage constrains language performance, whereas gradient analyses quantify how surviving regions are positioned within large-scale functional organisation. For example, temporoparietal hubs, damaged in many stroke survivors in our sample, typically occupy positions that coordinate interactions between control and default mode systems^72^; gradient analyses then capture how other language-relevant regions vary in their large-scale embedding following stroke and how this variation relates to behaviour. Importantly, variation in connectivity along a single gradient axis can show opposing associations with different language components, indicating that the behavioural consequences of reorganisation cannot be reduced to a simple gain or loss of function. Instead, the impact of altered large-scale coupling depends on the specific cognitive demands being considered. This framework situates behavioural variability within a low-dimensional functional architecture, showing how local damage and distributed changes in network embedding jointly shape language outcomes after stroke.

## 6. Future directions

While this study focused on language function, gradient-based functional organisation is likely to influence other cognitive domains, including executive function, memory, and spatial processing. Richer neuropsychological assessments of these functions would provide greater clarity about the cognitive consequences of gradient changes following stroke. Longitudinal studies are needed to track how connectivity gradients evolve across different stages of recovery, and how compensatory mechanisms develop in relation to structural damage. In addition, spared regions, often unaffected by MCA strokes, for example in anterior prefrontal regions, may play a critical role in supporting recovery through gradient-mediated network reorganisation. Understanding these dynamics could inform targeted therapies designed to boost neuroplasticity in chronic stroke.

## 7. Conclusion

Our findings demonstrate that gradient-based functional connectivity changes are associated with language outcomes after chronic stroke. Lesion–symptom mapping identified both classical perisylvian language regions and additional parietal, temporal, and occipital areas as critical for verbal working memory, comprehension, and writing. Connectivity gradient analyses revealed how surviving regions shift along DMN–control axes, capturing associations with both better performance (speech production via left IFG integration with DMN) and impaired performance (poorer writing following the same connectivity changes). Together, these complementary approaches provide a framework for understanding how focal lesions co-occur with distributed network dynamics, offering a new method for understanding the effects of stroke on behaviour.

## Credit authorship contribution statement

Ramya Balakrishnan: Conceptualization, Methodology, Formal analysis, Investigation, software, Writing—Original Draft, Writing—Review & Editing, Visualisation. Tirso Rene del Jesus Gonzalez Alam: Methodology, Software, Writing—Review & Editing. Robert Leech: Conceptualization, Methodology, Software, Writing—Review & Editing. Cathy J Price: Methodology, Investigation, Writing—Review & Editing. Elizabeth Jefferies: Conceptualization, Methodology, Investigation, Writing—Review & Editing, Supervision.

## Acknowledgments

We thank the past and present PLORAS team members, participants and volunteers who helped to collect data for this repository. Details of the PLORAS team members who collected the data can be found here: https://www.ucl.ac.uk/brain-sciences/ploras/project-information/team.

## Data availability

The PLORAS dataset used in this study can be made available to readers upon reasonable request.

## Open access

The authors have applied a CC BY public copyright licence to any Author Accepted Manuscript version arising from this submission.

