## Supplementary material for "Reorganisation of Functional Connectivity Gradients in Post-Stroke Aphasia"

### Section I: PCA Results for the Comprehensive Aphasia Test

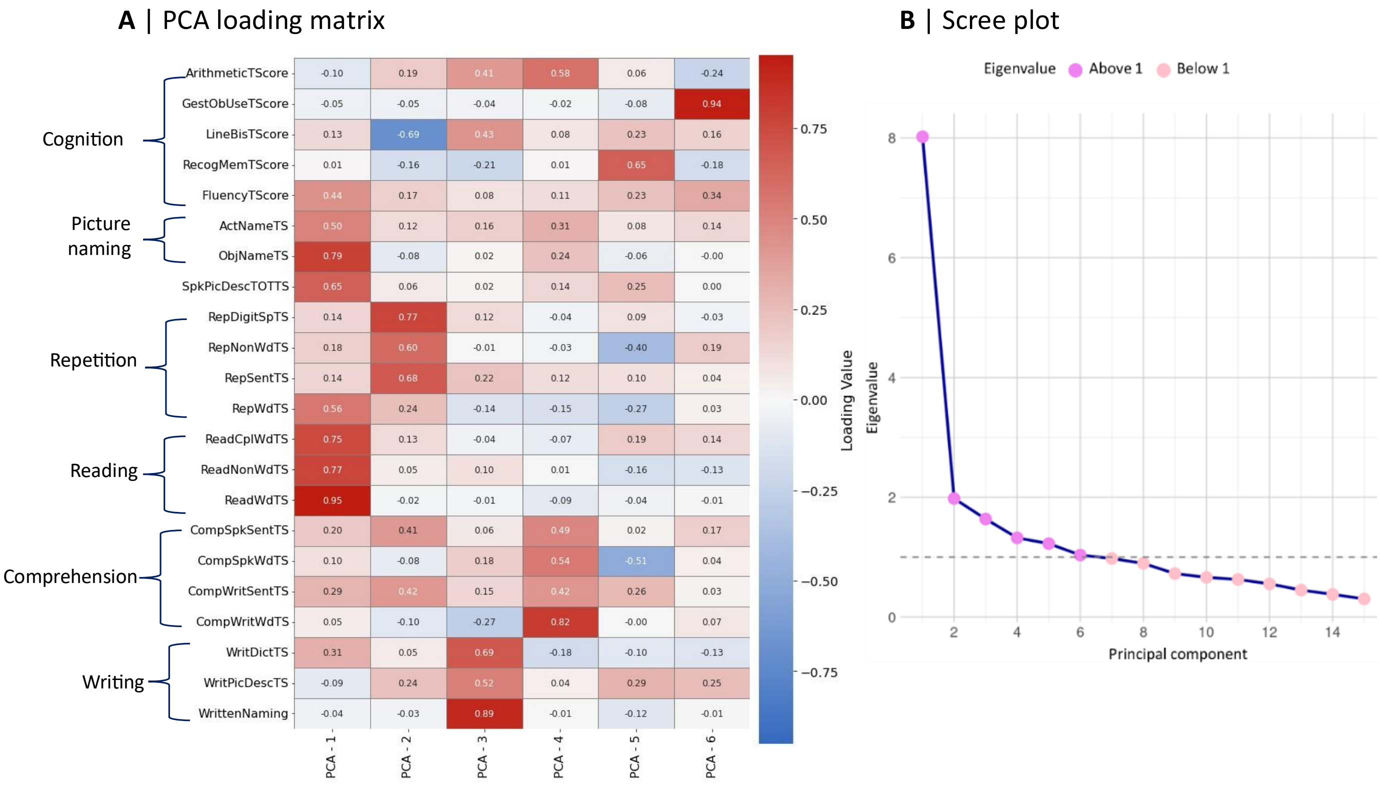


**Fig. 1: Principal component analysis results of the CAT. A)** PCA loading matrix of CAT subtests. PCA-1 is associated with speech; PCA-2 with phonology and working memory; PCA-3 with writing and visuomotor capacity; PCA-4 with comprehension and arithmetic; PCA-5 with recognition memory; and PCA-6 with gesture–object use. **B)** Scree plot showing the eigenvalues for each principal component.

### Section II: Post stroke difference gradients

We subtracted each stroke patient's connectivity gradient from the average gradient map of age-matched controls to focus only on stroke-specific effects, generating post-stroke difference maps for all three gradients. Figure 2 shows the average post-stroke difference maps for all three gradients (Fig. 2A–C). Positive values (red–yellow) in the difference maps indicate parcels with lower gradient values in patients than in age-matched controls, whereas negative values (green–blue) indicate parcels with higher gradient values in patients compared to controls.

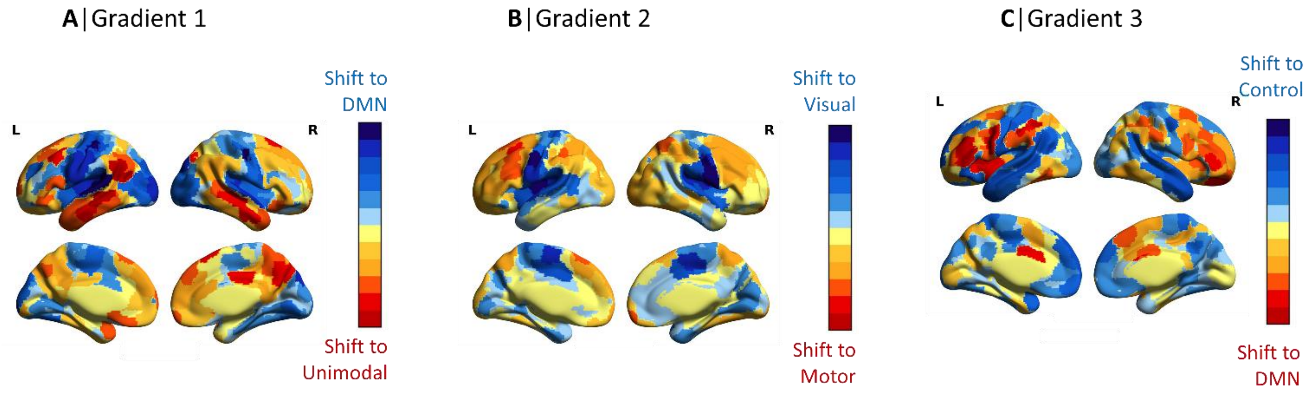


Fig.2: Post-stroke functional connectivity differences after removing aging effects. Average connectivity maps from age-matched controls were subtracted from each stroke patient's connectivity map, revealing positional shifts of brain regions across gradients. Top panel: A) shows stroke-related connectivity changes in Gradient 1, with shifts in both default mode network (DMN) and unimodal regions. B) shows connectivity changes in Gradient 2, involving shifts between visual and motor regions. C) shows connectivity changes in Gradient 3, with positional shifts in DMN regions and the control network. Warm colours indicate areas where patients had lower values than controls, while cool colours indicate areas where patients had higher values.

**Section III: Association between post stroke functional connectivity gradient changes and language outcomes within the language network**

We explored how post-stroke changes in functional connectivity gradients within the language network were associated with PCA-derived language outcomes. To focus on intact tissue, voxels with ≥25% lesion involvement were excluded from the language network mask (Fig. 3A). Results for PCA components 1-4 are in the main text. We additionally found worse performance in gesture–object use (PCA component 6) was associated with gradient changes in the left superior frontal gyrus, paracingulate gyrus, and juxtapositional lobule cortex (Fig. 3B). These regions, normally anchored toward the control end of Gradient 3, showed a shift in functional alignment toward the DMN, and this shift was linked to poorer gesture–object use (Fig. 3C).

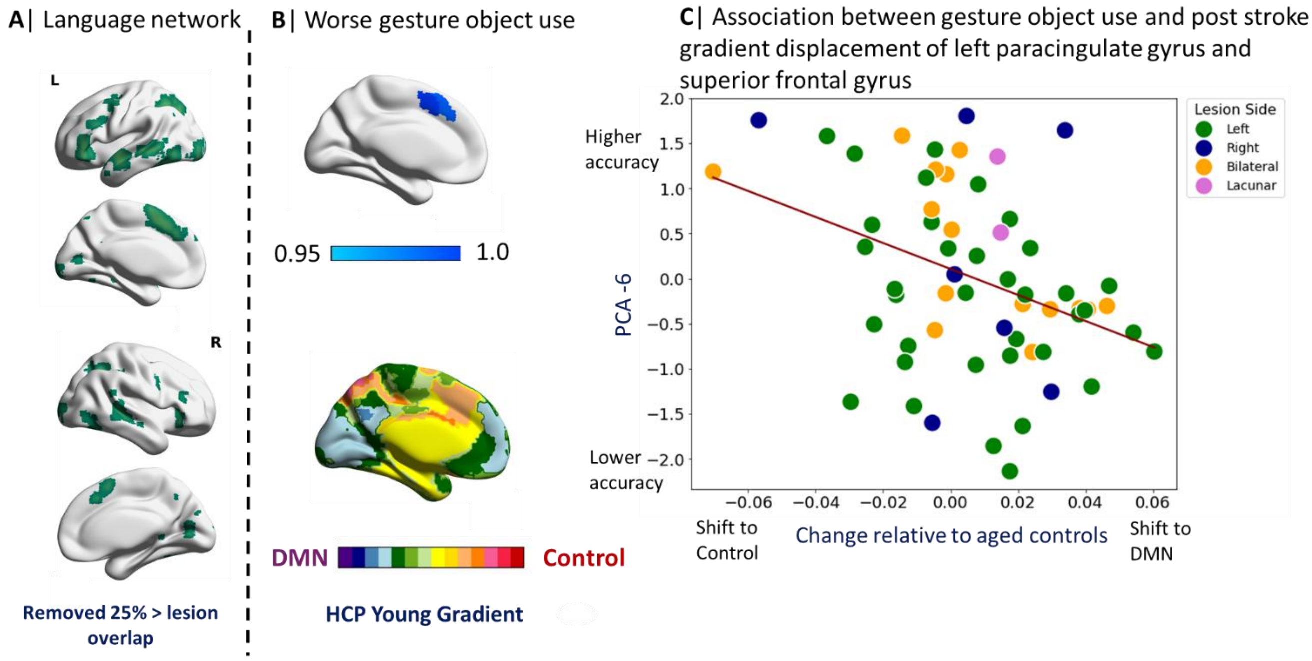


Fig. 3: Permutation test results within the language ROI. A) Language network mask used after removing lesion voxels for the ROI analyses. B) Poorer gesture–object use was associated with the left paracingulate and superior frontal gyri, regions normally aligned with the control-network end of Gradient 3. C) Scatterplot showing the relationship between shifts in these regions along Gradient 3 and gesture–object use performance.

**Section IV: Whole brain associations between post stroke functional connectivity gradient changes and language outcomes**

We investigated the association between changes in functional connectivity gradients in aphasia patients and PCA-derived language components by conducting lesion–gradient symptom mapping within largely intact brain regions (removing voxels lesioned in ≥25%) across the whole brain, to reduce the risk of type II errors. Nonparametric two-sample t-tests, using FSL's Randomise, with 5000 permutations were performed. Gradient 3 in whole-brain analysis showed clusters for both PCA component 5 (recognition memory) and component 6 (gesture–object use). Right frontal medial cortex, paracingulate cortex, and subcallosal cortex were associated with worse recognition memory (Fig. 4A) while the bilateral anterior cingulate cortex and juxtapositional cortex were associated with worse gesture object use (Fig. 4C). We further examined these regions' functional affiliation with the default mode network (DMN) and control network end: to visualise these associations, we plotted gradient values for these clusters from the post stroke difference gradient maps against task performance. Figure 4B shows that in aphasia patients, a shift of the right frontal medial, paracingulate, and subcallosal cortices toward the default mode network (DMN) was associated with poorer recognition memory. These regions normally occupy an intermediate position along Gradient 3. A similar pattern was observed in the anterior cingulate and juxtapositional lobule cortices, which are typically located nearer the control-network end of this gradient. In stroke patients, these areas also shifted toward the DMN pole, and this shift was linked to poorer performance on gesture object use (Fig. 4D).

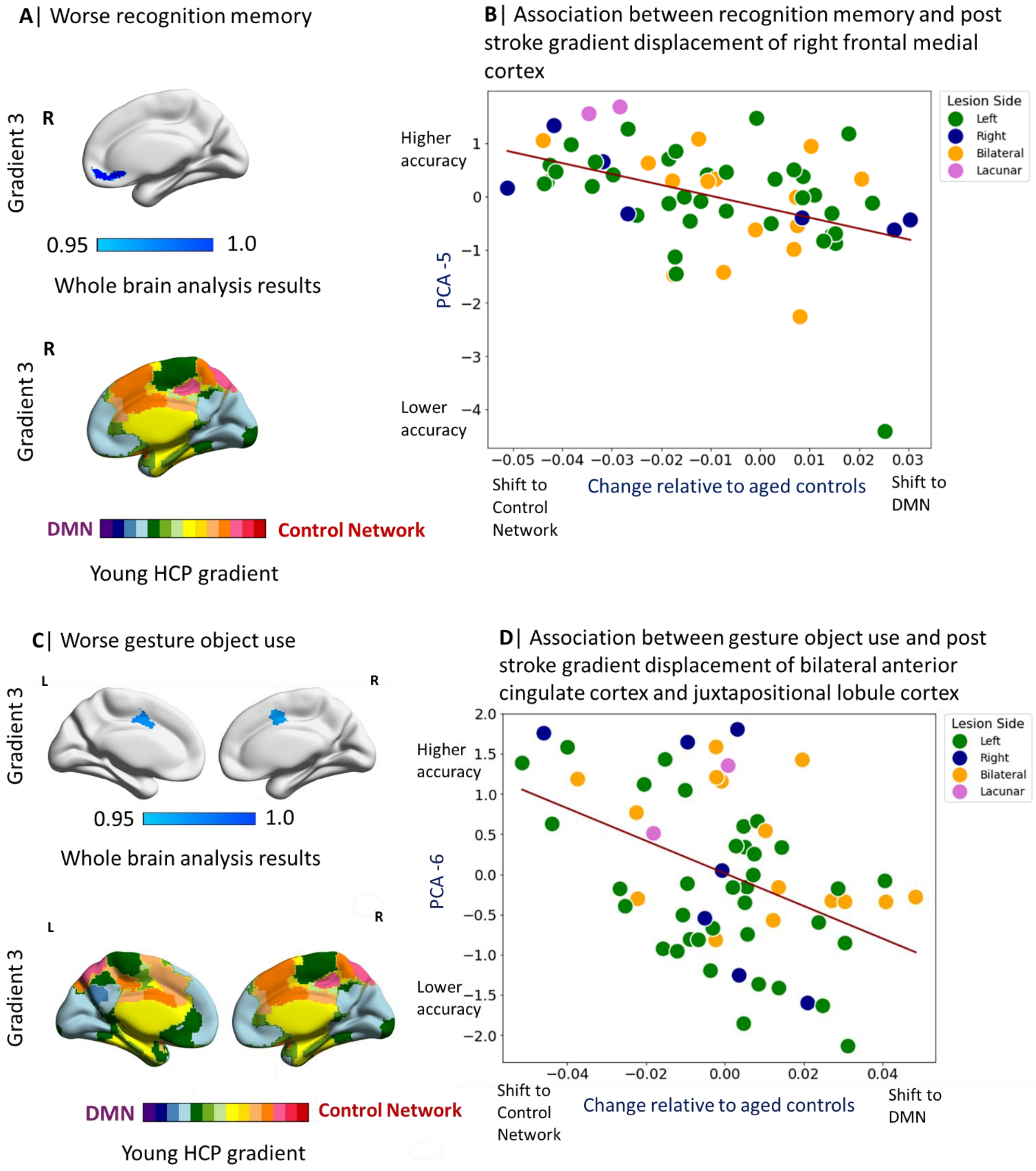

Fig. 4: Permutation test results at the whole-brain level. A) A significant cluster in the right frontal medial cortex was associated with poorer recognition memory performance (top left panel) along Gradient 3. This region normally occupies an intermediate position on this gradient. B) Scatterplot showing the association between the right frontal medial cortex's position along Gradient 3 and recognition memory performance (top right panel). C) Poorer gesture–object use was associated with bilateral anterior cingulate and juxtapositional lobule cortices (bottom left panel), regions typically located toward the control-network end of Gradient 3. D) Scatterplot showing the relationship between shifts in these bilateral regions along Gradient 3 and gesture–object use performance (bottom right panel).
